# *In vivo* x-ray diffraction and simultaneous EMG reveal the time course of myofilament lattice dilation and filament stretch

**DOI:** 10.1101/2020.03.03.970178

**Authors:** SA Malingen, AM Asencio, JA Cass, W Ma, TC Irving, TL Daniel

## Abstract

Muscle’s function within an organism depends on the feedback between molecular to meter-scale processes. While the motions of muscle’s contractile machinery are well described in isolated preparations, only a handful of experiments have documented the kinematics of the lattice occurring when multi-scale interactions are fully intact. We used time-resolved x-ray diffraction to record the kinematics of the myofilament lattice within a normal operating context: the tethered flight of *Manduca sexta*. Since the primary flight muscles of *Manduca sexta* are synchronous, we used these results to reveal the timing of *in vivo* cross-bridge recruitment, which occurred 24 (s.d. 26) ms following activation. In addition, the thick filaments stretched an average of 0.75 (s.d. 0.32)% and thin filaments stretched 1.11 (s.d. 0.65)%. In contrast to other *in vivo* preparations, lattice spacing changed an average of 2.72 (s.d. 1.47)%. Lattice dilation of this magnitude significantly impacts shortening velocity and force generation, and filament stretching tunes force generation. While kinematics were consistent within individual trials, there was extensive variation between trials. Using a mechanism-free machine learning model we searched for patterns within and across trials. While lattice kinematics were predictable within trials, the model could not create predictions across trials. This indicates that the variability we see across trials may be explained by latent variables occurring in this naturally functioning system. The diverse kinematic combinations we documented mirror muscle’s adaptability and may facilitate its robust function in unpredictable conditions.

## INTRODUCTION

Using ubiquitous molecular machinery, muscle performs diverse functions within an organism; at turns functioning as a motor, structural support, a repository for elastic energy, or as a shock absorber (Dickinson et al., 2000). A muscle’s functional output – the force it creates and its length change – depends upon multiscale interactions. Arrays of molecular motors interact (generating piconewton-scale forces) in a feedback loop with interacting muscle groups (generating newton-scale forces) within the animal’s body. The highly organized lattice of contractile machinery which powers contraction also changes shape as a result of internal forces, temperature, and externally applied forces. In turn, the shape of the lattice tunes force production.

For instance, increasing the spacing between molecular motors and the thin filaments to which they bind decreases binding probability, and therefore decreases force output (Williams et al., 2013). The spacing of the lattice co-varies with naturally occurring temperature gradients, and has the potential to shift the function of otherwise identical muscle sub-units from motors to springs and dampers (George et al., 2013; 2012). Additionally, the thick and thin filaments which house myosin molecular motors and actin binding sites are far from a rigid system; instead they stretch in conjunction with internal axial forces and muscle activation. Filament stretching has important mechanical implications, accounting for about 70% of muscle’s total compliance (Wakabayashi et al., 1994), and resulting in increased binding probability due to changes in the axial register of molecular motors and prospective binding sites (Daniel et al., 1998). While lattice dilation (Williams et al., 2013; Fukuda et al., 2005; Metzger and Moss, 1987) and filament stretch (Squire, 1997) have noteworthy mechanical implications, the relationship of their magnitude and timing within a naturally functioning organism has remained enigmatic. To address this, we documented molecular motor recruitment following nervous activation, filament stretching and changes in lattice spacing in a novel *in vivo* experimental system: the synchronous flight muscles of the hawkmoth *Manduca sexta*.

The advent of high-speed digital detectors in the early 2000’s made *in vivo*, time-resolved x-ray diffraction possible. Landmark studies of lattice kinematics in naturally functioning systems have largely focused on asynchronous insect flight muscles (fruitfly: *Drosophila* and bumblebee: *Bombus*) (Dickinson et al., 2005; Irving and Maughan, 2000; Iwamoto and Yagi, 2013). Irving & Maughan, for example, were the first to document the molecular kinematics of a mutant fly *in vivo*, connecting molecular mutations with their functional outcomes across the scales of animal flight behavior (Irving and Maughan, 2000). However one limitation of these systems is that in asynchronous muscle neural activation is decoupled from cycles of muscle shortening and lengthening; instead activation serves to keep the muscle in a contractile state by continuously suffusing myofibrils with calcium ions. Asynchronous muscle is uniquely specialized to power high-frequency wing flapping (Syme and Josephson, 2002). Iwamoto and Yagi leveraged this to record the mechanism of stretch activation independent of calcium release and re-uptake (Iwamoto and Yagi, 2013).

In contrast to asynchronous muscle, many mammalian skeletal muscles are activated in partial tetany, with multiple neural impulses stimulating larger contractile forces. Vertebrate cardiac muscle, on the other hand, has a one-to-one relationship between activation and contraction. In a manner analogous to cardiac muscle, the dominant flight muscles of *Manduca sexta* are synchronous with, generally, a one-to-one correspondence between neural activation and muscle contraction. Additionally, they mirror cardiac muscle in their function, contracting against fluid loads and acting predominantly on the ascending portion of their length-tension curve (Tu and Daniel, 2004).

**Fig. 1.**
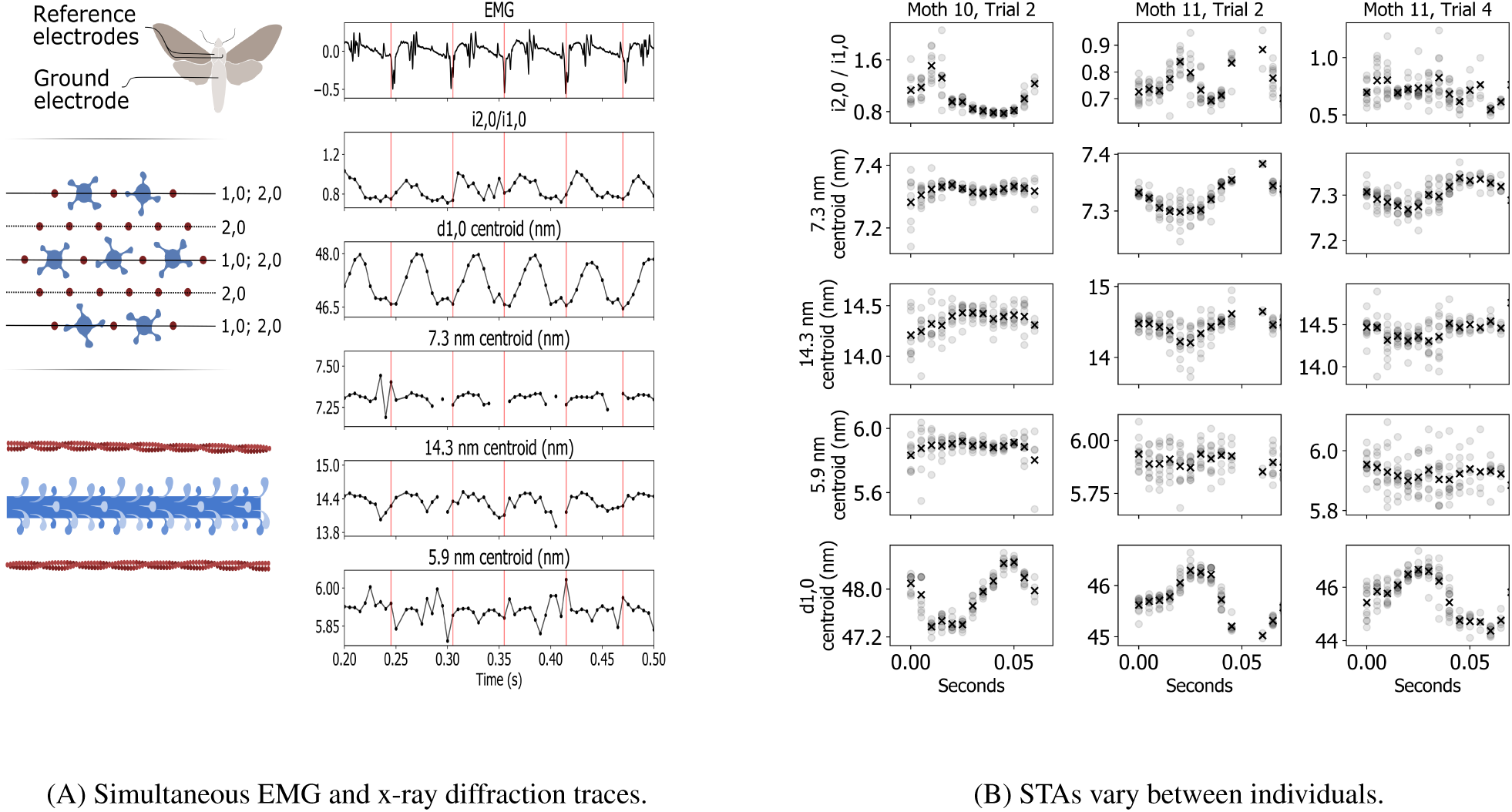
Time resolved lattice kinematics are consistent within a trial but vary between trials. (A) The muscle’s activation recorded in the EMG signal is indicated by a vertical red line. The time of the EMG peak was rounded to the nearest 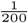 th of a second to match the frame rate of the detector, leading to a slight mismatch with the EMG recording which had a resolution of 25,000 Hz (Methods: Electromyogram). In the EMG trace shown, the large amplitude peaks are from the activation of the DLMs, while the EMG also picked up the activation of the neighboring antagonistic muscle group, the Dorsal Ventral Muscles. These are seen in smaller amplitude spikes. Cross-bridge recruitment was inferred by the radial shift of mass between the 1,0 and 2,0 planes as cross-bridges move towards the thin filaments following activation. Lattice spacing was measured using the 1,0 reflection. Thick filament stretching was inferred from changes in the spacing of the 7.3 nm meridional reflection and thin filament stretching from changes in the axial spacing of the 5.9 nm off-meridional reflection. (B) Data was phase averaged using the muscle’s endogenous depolarization. The mean of each time point following activation is marked with an X. All of the data we collected is shown in the supplement (Fig: S4).

By pairing simultaneous recording of muscle activation with x-ray diffraction data, we were able to phase average lattice kinematics using the organism’s natural activation. These results uniquely show the timing of molecular motor recruitment and the resulting lattice kinematics (stretching and dilation) following activation. Using phase averaged data we found that the molecular kinematics are consistent within individuals, often showing large excursions: the lattice can dilate by as much as 2.5 nm (5.4%), thick filaments can stretch by as much as 1.2%, and thin filaments by as much as 2.1%. In light of experimental and computational studies, the magnitude of these results support the notion that lattice kinematics significantly modulate muscle function (Daniel et al., 1998; Williams et al., 2013; Metzger and Moss, 1987). As is expected in a fully intact, normally functioning animal, there was considerable variation in both the timing and extent of lattice motions across individuals. These results highlight that while individuals may manifest the same functional outcome at the organism scale (flight), the mechanics of the lattice underneath can vary. Considering this, lattice kinematic data need to be considered within the context of individuals without supposing that latent variables are equal across individuals (Gomez-Marin and Ghazanfar, 2019).

## MATERIALS AND METHODS

### Animal preparation

To record cross-bridge recruitment and lattice kinematics as a function of electrical activation we used an insect synchronous muscle. The dorsal longitudinal muscles (DLMs) of *Manduca sexta* are synchronously activated, and are an excellent system for x-ray diffraction experiments since the thorax is composed of about a cubic centimeter of highly ordered muscle and the exoskeleton produces negligible scattering. The clear diffraction patterns produced by this muscle made it possible to collect time resolved data without frame averaging. We used a random mix of male and female moths 1-2 weeks post eclosion. We did not record sex and our limited sample size precludes analysis of sex differences.

We cold anesthetized a moth, tethered it and placed it in the beam line (Fig. 2). Moths were unencumbered except by Electromyography (EMG) electrodes inserted into the thorax and the ventral tether. The tether was a flattened stainless steel needle coated in cyanoacrylate glue inserted between the second and third coxae and crystalized with sodium bicarbonate. The moth was positioned on the tether with a pitch angle of approximately 30° to the horizon, similar to natural flight orientation. The moth was placed on the beamline with its body axis, and hence the axis of the dorsal longitudinal muscles, at a right angle to the beam’s incidence. The beam passed through the anterior, dorsal quadrant of the thorax, intersecting at approximately the d-c DLM subgroup based on external morphology (Eaton et al., 1988). It was aimed consistently in a small section where the wings would not obstruct the beam path. A hot air soldering iron set to its lowest heat setting was placed approximately 1m in front of the moth while a fan with a filter attached was placed behind the moth to collect dislodged scales. The warmth of the soldering iron and the wind current created by it and the fan stimulated natural flight behavior and kept the moth warmer than the room’s cool ambient temperature.

**Fig. 2.**
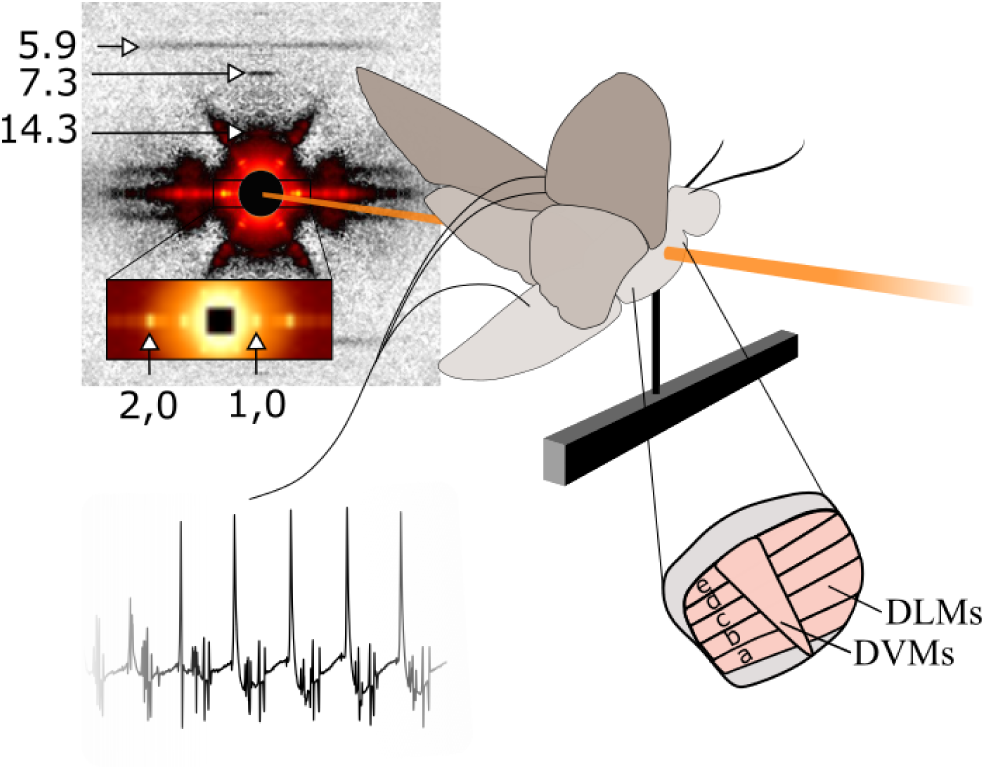
The moth was tethered within the beamline such that the beam passed through the dorsal-anterior portion of the thorax where the DLM’s comprise the bulk of the tissue volume. Simultaneous EMG data was collected to measure membrane depolarization. The x-ray diffraction image shown is after convex hull background subtraction, which leaves the reflections due to periodic structures intact, but removes the decaying intensity around the backstop due to the beam’s dispersion. This diffraction image is shown in isolation in the Supplement (Fig: S2).

### Electromyogram (EMG)

EMG electrodes with hooked tips were inserted into the posterior of the thorax after puncturing the exoskeleton with a needle. The ground electrode was inserted into the abdomen (Fig. 2). In order to record the rapid electrical transient of muscle activation the EMG must be recorded at high temporal resolution. Therefore we recorded this data at 25,000 Hz. Since the purpose of recording muscle activation was to correlate it with the kinematics measured using x-ray diffraction we rounded the timing of muscle activation to the nearest 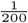 th of a second in order to correspond with the time base we used to record lattice kinematics. Because of this it may appear that there is jitter in the peak detection. The data (both x-ray diffraction and EMG) from animals that did not have periodic EMG signals with identifiable peaks for the full one second recording was excluded.

### X-ray diffraction

#### Beamline setup

The experiment was performed at the Biophysics Collaborative Access Team (BioCAT), beamline 18ID at the Advanced Photon Source, Argonne National Laboratory, Lemont, IL, USA (Fischetti et al., 2004). The beam energy was 12.0 keV with an incident flux of 10^13^ photons/s and attenuated as needed to 10^12^ photons s^−1^. It was focused to 250×250 *µ*m at the animal’s thorax, and 60×150 *µ*m at the detector with a sample to detector distance of 2 m. The sample was oscillated over a 1 mm excursion in the beam at 20 − 30 mm·s^−1^ in order to mitigate radiation damage. This method of oscillating the tissue relies upon the supposition that all sarcomeres in a local region are behaving in the same manner, so moving the sample will not alter the x-ray periodicities observed. We used a photon counting Pilatus 3 1M (Dectris Inc.) with 981×1043, 172×172 *µ*m, pixels with a 20-bit dynamic range. Raw image data was collected at 200 Hz with 1 ms of dead time per frame, and with the X-ray shutter open continuously.

#### Data reduction from x-ray diffraction images

The 32-bit tagged image file format (TIFF) image stacks returned by the x-ray detector were annotated using the Musclex software suite developed by the Biophysics Collaborative Access Team (BioCAT) (Jiratrakanvong et al., 2018). Images were annotated by two different individuals and cross-validated to ensure that there was not a substantial difference between them (Fig: S3). Trials where data was not traceable were excluded. Each image was quadrant folded to center and average intensities across axes of symmetry and annotated for meridional and equatorial intensity peaks. We used versions of Musclex prior to version 1.14.12. These have an error in the image centering algorithm that results in the image center being rounded to the nearest pixel. This means there is a center placement error of <0.5 pixels that could result in a maximum radial compression of the image of 0.5 pixels. An improvement in centering is being implemented in future versions which reduces the error inherent in interpreting physically continuous data recorded in a pixel framework by remapping intensity based on the calculated image center.

#### Cross-bridge recruitment

cross-bridge recruitment can be inferred by the shifting of mass away from the thick filaments and towards the thin filaments. Since the flight muscle of *Manduca sexta* has a packing ratio of 1:3 thick to thin filaments, the intensity of the 1,0 peak contains thick and thin filament mass, while the 2,0 peak is the sum of the second-order harmonic of the 1,0 and the intensity due to layers containing only thin filaments (Fig: 1). As cross-bridges move towards the thin filaments, there is a decrease of the 1,0 intensity due to the shifting of mass towards the layers of thin filaments. In contrast, the 2,0 stays relatively constant since there is little change in the total mass on these planes. Therefore the ratio of 2,0 to 1,0 intensity is expected to be highest during peak cross-bridge binding. This differs from vertebrate muscle, where the ratio of the 1,1 reflection and 1,0 reflection is a surrogate for cross-bridge binding. The intensities of the 1,0 and 2,0 reflections were computed using Musclex’s “Equator” function, which first sets a box around the equator, uses convex hull background subtraction, and finally calculates peak intensity.

#### Lattice spacing

Lattice spacing was measured by tracking the 1,0 reflection’s centroid motion using Musclex’s Equatorial function and a Voigt model fitted to three peaks: 1,0; 1,1 and 2,0. This data was gathered simultaneously with cross-bridge recruitment. Notably, the Voigt model we used within the Equator function’s framework is over-determined with only two peaks, so we tracked the 1,1 reflection intensity in the fit model as well. The distance of the centroid of the 1,0 equatorial reflection from the backstop correlates by Bragg’s law to the distance between layers of the filament lattice containing the thick filament.

#### Filament strain

Thick and thin filament strain were calculated using axial filament spacing changes recorded along the meridian and in layer lines of the diffraction pattern. We used Musclex’s Diffraction Centroids function to estimate the axial spacing of the following intensity peaks: the 14.3 nm meridional, an indicator of spacing changes between layers of myosin crowns; the 7.3 nm meridional, an indicator of spacing changes in the myosin containing thick filament backbone; and the 5.9 and 5.1 nm off meridional actin layer lines, which correspond to the pitches of the left and right genetic helices. At the exposure times we used, the actin layer lines were relatively weak. The 5.9 nm layer line intermittently yielded reliable data, while the 5.1 nm layer line was not measurable for most trials. The data presented for the 5.9 nm layer line should be interpreted with due caution since without 5.1 nm data filament twisting and stretching cannot be differentiated.

### Data handling

All of the code we used can be found in the GitHub repository: In_vivo_Myofilament_Lattice_Kinematics. All of the data that we used in preparing this manuscript can be found on Dryad at the link: https://doi.org/10.5061/dryad.s1rn8pk51 under the same name as this manuscript.

#### Permutation bootstrap

To assess if a signal contained significant cyclic changes at the same frequency as the muscle’s activation we used a permutation bootstrap. We took the Fourier transform of a time series signal, which returned the power of the signal across frequency space. We then compared the power of the raw signal at wing beat frequency to the power obtained for a signal composed of the original signal, but randomly shuffled. If out of 1,000 permutations, fewer than 5% had a power greater than or equal to the original signal’s power at wing beat frequency the signal was used to compute average periodic excursions. If more than 5% had a power greater than or equal to the original signal’s, we cannot evaluate what component of the data is noise, so the signal was excluded from computing excursions. Out of the 11 trials we recorded, 11 showed significant periodic changes in lattice spacing, 9 showed significant periodic changes in cross-bridge recruitment, 7 showed significant periodic thick filament stretch, 6 showed significant periodic changes in the 14.3 nm centroid spacing and 6 showed significant periodic thin filament stretch. These results correspond to the annotator’s observations that data encoded on the equator is typically much stronger, while at the exposures lengths we used data encoded along the meridian was more challenging to track.

#### Spike triggered average (STA)

The STA is created by phase averaging time course x-ray diffraction data using the muscle’s depolarization as the start point. One phase is defined to be the period of time between sequential membrane depolarizations (the interspike interval). Interspike intervals were denoted using Python scripts which identify EMG peaks and down sample their time resolution to that of the x-ray data, indicating the frame during which depolarization occurred.

#### Correlation

The correlation of signals as a function of proportion of interspike interval lag was used to determine if there was a consistent phase offset of cross-bridge recruitment and thick filament stretch, and of cross-bridge recruitment and lattice spacing change across trials. We also calculated the correlation of two signals as a function of absolute time lag (Fig: S5) and found no consistent pattern. Correlation was calculated using Numpy’s built in correlation function.

#### Gradient boosted decision tree mechanism-free model(xGBoost)

xGBoost is a library to create gradient boosted decision trees that can be used for regression, classification and prediction (Chen and Guestrin, 2016). Gradient boosting iteratively combines weak hypotheses (in the case of xGBoost a weak hypothesis is a decision tree that has an imperfect prediction) that follow the gradient descent of an error loss function. At each iteration, a new tree is added with the goal of fixing the error left in the model prediction from the previous iterations. The loss function can be generalized as any differentiable function. xGBoost is a particularly fast and effective implementation of the machine learning algorithm ‘gradient boosted decision trees’ that has been used to win many Kaggle competitions (Cook, 2020). We used it to create a mechanism-free prediction of lattice dilation from filament stretching and cross-bridge recruitment data. Specifically, the model used the 14.3 nm reflection’s intensity (often used as an indicator of cross-bridge recruitment in vertebrate systems), the ratio of the reflections of the 2,0 and 1,0 planes, the 7.3 nm reflection’s centroid and the 5.9 nm off meridional’s centroid. Since the change of the lattice’s shape from one timestep to the next depends not only on the state of these parameters, but also their time history, we extended the information available to the model by including the time history of cross-bridge recruitment and filament stretching up to 11 time steps (0.055 s) prior to the time point to be predicted.

The xGBoost model is instantiated using hyperparameters. Optimizing the choice of hyperparameters for the model to create predictions for a given dataset can be handled by methods like grid search, coordinated descent, or the genetic algorithm. We chose to use a genetic algorithm. In essence, a small set of models (10 in our case) are constructed with randomly selected hyperparameters; these are a first generation. The next generation is created by combining the hyperparameters of the most successful models in the parent generation, along with random “mutation” to one of the hyperparameters. We used a total of 7 generations and allowed the crossing of 4 of the parameter models. We adapted code for an implementation of the genetic algorithm for selecting hyperparameters for xGBoost that can be found on the GitHub repo “Hyperparameter-tuning-in-XGBoost-using-genetic-algorithm” (Jain, 2018*a*;*b*) In the interest of reproducibility, the model’s hyperparameters can be found in the supplement (Table S1,2; Supplement: xGBoost).

## RESULTS

We recorded cross-bridge recruitment, axial filament strains, and radial lattice dilation as a function of endogenous muscle activation during *in vivo* tethered flight. These were indicated respectively by the ratio of the intensities of the 2,0 to the 1,0 equatorial reflections; axial shifts of the centroid of the 7.3 nm meridional reflection and axial shifts of the 5.9 nm actin off meridional reflection, and finally radial shifts in the position of the equatorial reflections (Fig: 1A). We also recorded the axial shifting of the 14.3 nm meridional reflection. With a detector frame rate of 200 Hz and wing beat frequencies ranging between 13-19 Hz (mean 16 (s.d. 2), we obtained 11-15 x-ray diffraction images from each cycle of shortening and lengthening. Each trial lasted one second, and we created spike triggered averages (STAs) by phase averaging the data based on the peaks of each individual’s electromyogram (EMG). Most individuals had consistent STAs, however there was variability between individuals in the time course and excursions of their STAs for a given data type (Figs: 1, S4). We calculated the excursion of a signal as the difference between the maximum and minimum of the STA for each trial.

### The timing and extent of cross-bridge recruitment, filament stretching, and lattice dilation are revealed in spike triggered averages (STAs)

By using a synchronous muscle group, we were able to correlate muscle activation and the resulting recruitment of myosin molecular motors to the thin filaments. The peak cross-bridge recruitment occurred an average of 0.024 (s.d. 0.026) seconds following activation, with a resolution of 0.005 seconds. We measured the extent of filament stretching by calculating the excursion (maximum minus minimum) of the 7.3 nm reflection’s STA. This revealed that thick filaments stretch by an average 0.75 (s.d. 0.32)% across 7 trials. By first calculating the excursion of each trial and subsequently computing the mean across trials we avoid the blunting of the signal that can occur if the excursion is calculated from the amalgamation of all trial’s normalized STAs. Likewise, we calculated the lattice’s dilation as the maximum minus the minimum of the STA for each trial, with a mean taken across all 11 trials. The lattice dilated by 2.72 (s.d. 1.47)%, which corresponds to 1.24 (s.d. 0.66) nm.

For each of these data types, the pattern of the STA was relatively consistent across many cycles of shortening and lengthening within an individual trial. This is especially apparent in the case of the d1,0, which, as the clearest diffraction signal, is least subject to error during annotation. However the STAs reveal extensive variation across individuals, which means that there is a large standard deviation in the timing and extent of myofilament lattice kinematics.

### Interrelationship of lattice kinematics: inter- and intra-indiviudal patterns

Our kinematic data capture filament motions within a mechanically coupled system. Here we explore whether there are consistent relationships between the various lattice kinematics within individuals and, additionally, whether there are patterns common to all individuals. Since thick filament strain results from active tension development, passive tension development and filament activation (Wakabayashi et al., 1994; Ma et al., 2018; Irving et al., 2011; Piazzesi et al., 2018), we hypothesized that the stretching of the thick filament indicated by movement of the 7.3 nm meridional reflection would be maximally correlated with increased cross-bridge binding at a relatively fixed phase offset. However the maximum cross-correlation between cross-bridge recruitment and thick filament stretch shows variable timing offsets across individual trials, demonstrating a complicated relationship between these variables. This may be partially explained by recent work which demonstrates that there is a non-linear relationship between tension and thick filament extension (Ma et al., 2018). Proteins similar to titin may also contribute to sarcomere elasticity in *Manduca sexta* (Yuan et al., 2015) with significant non-linear behavior (Powers et al., 2018; Trinick, 1996; Tskhovrebova et al., 1997) and temperature dependence (Bullard et al., 2006). Finally, while stretching of the filaments indicates internal axial force, cross-bridges also exert force radially. This means that the magnitude of the component of cross-bridge force applied axially changes as a function of the angle between motors and binding sites (Schoenberg, 1980; Williams et al., 2010). In light of these considerations, it would be remarkable if the movement of mass towards the thin filaments alone explained the periodic stretching of the thick filaments, or the periodic dilation of the lattice. But taken together, can we use these data to fully model the system?

If the data are sufficient to explain one another mechanistically, even though the linkage is non-obvious, they should be sufficient to train a mechanism-free, data-based model. (Though by creating a mechanism-free model we cannot exclude the possibility that the data are insufficient to explain one another mechanistically, but contain enough correlative information to successfully predict one-another.) We used a gradient boosting decision tree algorithm housed in the xGBoost library (Chen and Guestrin, 2016) to train a mechanism-free model. First we trained the model with 75% of a trial’s data, using the other 25% as a test set to evaluate model performance. We set up the model to predict lattice spacing from filament stretching and cross-bridge recruitment since lattice spacing is typically the cleanest of the signals tracked in x-ray diffraction due to its intensity. In addition to using the current state of filament stretch and cross-bridge recruitment to predict the current lattice spacing, we also provided the model with the time history of the predictor variables up to 11 time-steps (0.055 s) previous to the current state. For these within-trial predictions the random forest-based model performed well, with a Root Mean Square Error (RMSE) of 0.27 nm, demonstrating that it is possible to predict lattice spacing change from filament stretching and cross-bridge recruitment within a trial.

We then addressed our larger question – can we create a prediction of lattice spacing change from cross-bridge recruitment and filament stretching data that holds across individuals? We iteratively withheld the data from one individual as a test case and trained the model with the data from all other individuals. The average RMSE across all trials was 0.78 nm. This shows that the themes the model uses to create intra-individual predictions do not generalize well across individuals.

**Table 1.** For each of the 11 total trials, and each data type we ran a permutation test to see if there was a significant frequency component at wing beat frequency. If the chance of a random permutation yielding a power as high or higher than the raw data at wing beat frequency was less than 5% we factored it into this summary. If it was greater than 5% we can only conclude that if there are periodic changes occurring, they are of lower amplitude than the noise envelope. The number of trials that passed the permutation test is recorded in the column “Number of trials”. For each trial and datatype, its baseline value was the minimum point in the STA. The average baseline as the mean of across all trials. The excursion of each datatype was calculated as the average across all trials of the STA maximum minus the STA minimum. The percent excursion was the absolute excursion divided by the baseline for each trial – these were averaged across all trials. Finally the peak time was the time elapsed from the beginning of the STA to its maximum, which was averaged across all trials.

| Data type | Number of trials | Baseline | Excursion | Percent excursion | Peak time |
| --- | --- | --- | --- | --- | --- |
| i2,0/i1,0 | 9 | 0.708 (s.d. 0.076) | 0.250 (s.d. 0.204) | 35.35 (s.d. 26.92)% | 0.024 (s.d. 0.026) s |
| 7.3 nm centroid | 7 | 7.29 (s.d. 0.02) nm | 0.06 (s.d. 0.02) nm | 0.75 (s.d. 0.32)% | 0.058 (s.d. 0.024) s |
| 5.9 nm centroid | 6 | 5.87 (s.d. 0.04) nm | 0.07 (s.d. 0.04) nm | 1.11 (s.d. 0.65)% | 0.044 (s.d. 0.025) s |
| d1,0 | 11 | 45.84 (s.d. 0.88) nm | 1.24 (s.d. 0.66) nm | 2.72 (s.d. 1.47)% | 0.032 (s.d. 0.018) s |
| 14.3 centroid | 6 | 14.26 (s.d. 0.08) nm | 0.30 (s.d. 0.13) nm | 2.08 (s.d. 0.91)% | 0.039 (s.d. 0.026) s |

## DISCUSSION

Combining high-speed, time-resolved x-ray diffraction with simultaneous recording of the electrical activation of the synchronous flight muscles of the hawkmoth *Manduca sexta* we resolved myofilament lattice kinematics during fully intact tethered flight. Taken together, our data reveal intra-individual patterns of axial myofilament stretching, radial spacing changes, and cross-bridge recruitment that follow the endogenous activation of muscles. This method gives us a window into the molecular motions that underlie muscle force production.

**Fig. 3.**
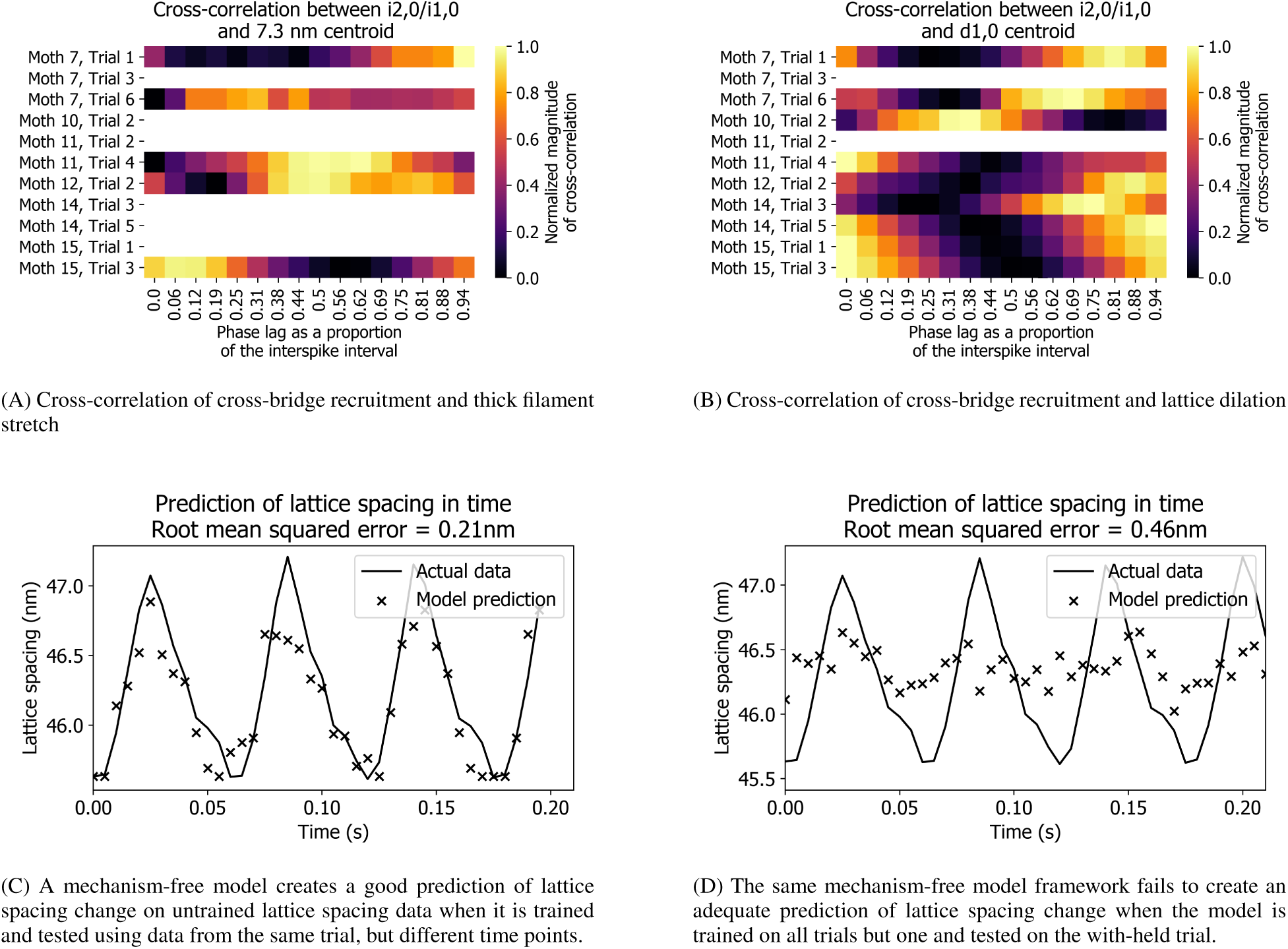
The relationship between lattice kinematics is not consistent across trials. Panel A) shows that there is not a consistent phase offset of cross-bridge recruitment and filament extension. The magnitude of the cross-correlation is normalized, and phase offset is measured in proportion of the interspike interval (the period of time between subsequent muscle activation). Some trials do not contain adequate data for all data pairings – a permutation bootstrap was used to determine for each datatype within a trial if it contained a significant periodic component at wingbeat frequency, and those that did were included. Correlation as a function of time lag in seconds can be found in the Supplement (Fig: S5). Panel B) shows the correlation between cross-bridge recruitment and lattice spacing. Panels C) and D) show a mechanism-free model’s prediction of lattice spacing from filament strains and cross-bridge recruitment in time. Panel C) shows a prediction for the case where the model is trained on 75% of a trial’s data, and tested on the other 25%. The average Root Mean Squared Error (RMSE) was 0.27 nm. Panel D) shows a representative image from the case where the model was trained on the data from all trials, except for one that was with-held as a test set. The average RMSE across all trials trained in this manner was 0.79 nm. The disparity between the model’s performance under these paradigms suggests that while this model is capable of making good predictions from these types of data sets (panel C), the predictions made by the model cannot extrapolate effectively across individuals for this data set (panel D).

### Cross-bridge recruitment

While x-ray evidence for the timing of the excitation-contraction pathway has been used in isolated fibers (Reconditi et al., 2011), the time course of cross-bridge recruitment following activation for fully intact and *in vivo* preparations has not previously been reported. Muscle contraction is triggered by motor neuron activation, leading to depolarization of the muscle cell membrane and the subsequent release of calcium from the sarcoplasmic reticulum into the cytoplasm surrounding the filament lattice. Calcium binding initiates the shifting of the troponin - tropomyosin complex away from actin binding sites and the dissociation of molecular motors from the thick filament, allowing the formation of cross-bridges between the filaments (Gordon et al., 2000; Squire and Morris, 1998; Stelzer et al., 2006). Our data show that peak cross-bridge recruitment occurs an average of 0.024 (s.d. 0.026) seconds following activation, to a resolution of 0.005 seconds. While the time course of cross-bridge recruitment was generally consistent in an individual trial over many cycles of sequential shortening and lengthening, there was variation between individuals, resulting in the large standard deviation.

The variation in lattice kinematics that we recorded in a fully intact system could arise from several latent variables. For example, temperature varies both spatially and temporally in *Manduca sexta*. Temperature gradients correlate with variation in lattice kinematics at the molecular-scale, and molecular-scale variation corresponds to functional gradients across the muscle group at the organism scale (George et al., 2013). In addition to gradients across a muscle group, the mean temperature of the muscle group increases with increasing wing beat frequency in *Manduca sexta* (Heinrich and Bartholomew, 1971). Temperature may contribute to the observed variation in our system since the diffusion of substrates like calcium ions and molecular motors is slower in cool muscle. However the experimental constraints of simultaneous EMG recordings and high speed, time-resolved x-ray diffraction in a naturally functioning animal limit our ability to resolve spatio-temporal patterns of temperature in the muscle. In favor of an intact preparation, we did not control temperature and all of the moths were flapping their wings at different frequencies of their own volition. While temperature likely contributes to the variation we observe in our data, diversity in the timing of maximum cross-bridge recruitment could also arise from other mechanisms, such as other steps in the excitation-contraction pathway, or co-occurring lattice spacing changes and filament stretching.

Although the ratio of the intensities of the 2,0 and 1,0 reflections is often used to quantify the shifting of mass between them (Irving, 2006), peak intensities could in theory be affected by changes in lattice ordering (Bershitsky et al., 2009) over a cycle of shortening and lengthening. Additionally, shifting of mass from the thick to the thin filaments does not necessarily imply binding. However, despite these caveats, the shifting of mass is well correlated with strong motor binding in vertebrate muscle (Squire, 1997; Harford and Squire, 1992). This ratio is an accessible surrogate for cross-bridge recruitment during time-resolved x-ray diffraction.

### Filament stretching

Filament stretching modulates muscle’s function. Axial filament stretching accounts for a large component of the sarcomere’s total compliance (about 70% (Wakabayashi et al., 1994)), enabling the return of stored elastic strain energy. The stretching of filaments also means that cross-bridges do not act independently of one another. Instead, as the relative separation between binding sites and cross-bridges changes, binding probability is also altered. Spatially explicit modeling demonstrates that these changes in axial register may mediate force output (Daniel et al., 1998). In addition to changes in axial register, strain in the thick filament may alter its twist. In *Lethocerus* asynchronous flight muscle thick filament strain is accompanied by a change in twist of about 12°that would help move heads closer to actin target zones as thick and thin filaments are strained (Perz-Edwards et al., 2011). Unfortunately X-ray patterns from *Manduca sexta*’s flight muscle lack the rich layer lines that *Lethocerus* muscle displays, so we could not identify changes in helical twist of the thick filament. The extent to which filament twist occurs in *Manduca sexta* as a function of filament extension remains conjectural until other experiments are devised to detect such changes.

Despite its functional significance, the extent of filament stretch occurring during natural muscle function was unknown until recent evidence from time-resolved x-ray diffraction revealed subtle (0.2 %) thick filament stretching in the dominant flight muscles of *Drosophila* (Dickinson et al., 2005). Our measurements show that the thick filament stretches by an average 0.75 (s.d. 0.32)% across 7 trials as indicated by changes in the 7.3 nm meridional intensity peak (Fig: 2). While the symmetry of *Manduca sexta*’s thick filament is not yet known it may be different from vertebrate filaments, possibly akin to *Lethocerus*, which has four-fold rotational symmetry (Hu et al., 2016; Reedy et al., 2000). Thus the 7.3 nm reflection may not be exactly equivalent to the M6 reflection in vertebrate muscle, nor the 14.3 nm reflection to the M3. With this caveat, in analogy with vertebrate muscle, the 7.3 nm reflection likely includes contributions from the second-order of the 14.3 nm myosin head repeat with additional contributions from the thick filament backbone (Brunello et al., 2006). The imperfect correlation of the spacing and intensity changes of the 7.3 nm reflection and the 14.3 nm reflection in vertebrate muscle, however, indicates that the 7.3 nm reflection is dominated by structures in the backbone; meanwhile, the 14.3 nm reflection is dominated by the periodicity of the myosin heads so that spacing changes in the 7.3 nm reflection may be used as a measure of thick filament extensibility (Brunello et al., 2006; Huxley et al., 1998; Linari et al., 2000; Ma et al., 2018). We have shown the cross-correlation of the 7.3 nm and 14.3 nm repeats from these data in the Supplement (Fig: S1). We conjecture that the same pattern seen in vertebrate muscle holds for the flight muscle of *Manduca sexta*. Though larger than those observed in *Drosophila*, the values we report are near those observed in isolated vertebrate fibers under isometric contraction (Wakabayashi et al., 1994; Brunello et al., 2006; Huxley et al., 1994). This result confirms the relevance of these parameters to *in vivo* function in a synchronous muscle group during natural function. Additionally, by measuring the extension of filaments within individuals rather than averaging across trials, the blunting of the signal that occurs when averaging phase offset data is avoided.

### Lattice dilation

Muscle function is also impacted by the spacing of the myofilament lattice, which is thought to play a role in the Frank-Starling mechanism in the mammalian heart (Moss and Fitzsimons, 2002). cross-bridge binding alters lattice spacing, and in turn their binding probability and the direction of the forces they generate are regulated by lattice spacing (Schoenberg, 1980). Radial cross-bridge extension has also been proposed as a site of elastic energy storage that could be returned to power cyclic contraction (Williams et al., 2012; George et al., 2013). Through mechanisms such as these, lattice spacing change results in a steeper length-tension curve than would be produced if only filament overlap changed during contraction (Williams et al., 2010). Since these muscles act on the steep ascending portion of their length-tension curve they generate larger forces in response to perturbations that stretch them, without a need to modify nervous control. Therefore in the case of cardiac muscle when blood pressure suddenly rises and increases ventricular filling, or in the case of flight muscle when a gust of wind buffets against the wing, the muscle autonomously contracts more forcefully (Tu and Daniel, 2004). This is an example of a cell-scale set point that yields a reflexive response to environmental perturbations at the organism scale, lending robustness to rapidly changing external demands (Moss and Fitzsimons, 2002).

Isolated muscle preparations have yielded mixed interpretations of how lattice spacing changes over the course of a contraction, although it is clear that lattice spacing is influenced by both mechanical and electrostatic interactions (Smith, 2014). Often the lattice is assumed to have either a constant spacing during contraction, or to be isovolumetric. But these assumptions don’t capture the extent of lattice spacing change observed experimentally. For instance, in skinned fiber preparations it was shown that during force generation lattice spacing tended toward a spacing near that observed when the membrane was intact (Millman, 1998). Meanwhile it was shown in relaxed, intact vertebrate muscle fibers that the sarcomere can be approximated as being isovolumetric, but when cross-bridges are active both axial tension and sarcomere length are determinants of lattice spacing (Bagni et al., 1994). Are lattice spacing changes occurring during *in vivo* muscle function? The average lattice spacing excursion we measured across 11 trials was 2.72 (s.d. 1.47)%, corresponding to 1.24 (s.d. 0.66) nm. These data stand in contrast to the results of Irving *et al* (Irving and Maughan, 2000), which showed no measurable lattice spacing change in the flight muscles of *Drosophila* to a resolution of 0.05 nm; the equivalent of *±*0.1% lattice spacing change in their system. While these are significant changes in spacing, they are smaller than the approximately 4% change in spacing needed to maintain a constant volume based upon the strains of this muscle group (about 9% (Tu and Daniel, 2004)). The average lattice spacing change we measured corresponds to an alteration in the force generated in osmotically compressed demembranated myofibrils of nearly 20% (Williams et al., 2013). Therefore we expect that lattice spacing change may be an important determinant of the force produced by this muscle group. Moreover, the maximum lattice spacing excursion we recorded was nearly twice the mean (2.5 nm). Akin to cross-bridge recruitment and filament stretching, lattice spacing change followed a stereotyped time course within individual trials, but the time course of lattice dilation demonstrated great breadth across trials. Similar parameters to those noted for cross-bridge recruitment and filament stretch may also give rise to the variation we observed in lattice spacing.

### Interrelationship of lattice kinematics: inter- and intra-indiviudal patterns

Ultimately, the myofilament lattice is a mechanically coupled system, but we cannot intuit how each of the pieces relate to one another across the widely variable STAs that we documented. Moreover, cross-correlation did not reveal thematic phase relationships between kinematics across trials, but rather highlighted the variation in the timing of kinematics relative to each other.

In response to these limitations, we built a mechanism-free machine-learning model to predict lattice dilation using the other kinematics we recorded. While the model performed well within trials, it was not able to forecast across trials effectively. The model cannot find relationships in the training set that explain the test set data, and instead of interpolating the model must extrapolate to forecast across trials, reflecting the visibly variable STAs. This may indicate that there are mechanisms that we did not record which account for the inter-trial variation. Latent variables like temperature, externally applied forces, the timing of activation, and antagonistic muscle activation are components that may be necessary to mechanistically explain myofilament lattice dynamics. These results demonstrate that the myofilament lattice uses a panoply of kinematic combinations, the breadth and significance of which we have yet to grapple with during natural function.

## CONCLUSION

At the organism scale muscle exhibits diverse functionality, at turns powering motion, stabilizing the body, storing energy, and dissipating energy (Dickinson et al., 2000). Molecular to organism scale feedback enables muscle to meet these demands. We recorded myofilament lattice kinematics during natural behavior at all scales, revealing that thick filaments stretch by 0.75 (SD: 0.32)% and that the lattice dilates by 2.72 (SD 1.47)%. By using the synchronous flight muscles of *Manduca sexta*, we were able to record that peak cross-bridge recruitment occurs 24 ms (SD: 26 ms) after activation *in vivo*. Despite the inherent uncertainty in interpreting x-ray diffraction data from a muscle group with unresolved ultrastructure, this system holds promise for understanding the *in vivo* dynamics of a muscle group with striking similarities to human cardiac muscle. We recorded extensive inter-individual variation in the timing and extent of lattice kinematics that could not be predicted by a powerful, mechanism-free model. The machine learning model we used capitalizes on weak patterns in data to make predictions, regardless of whether those patterns are mechanistic or correlative relationships. The inability of this model to forecast across trials indicates that latent variables (such as variation in the timing of muscle activation and the interaction of many muscles in the animal’s body) may give rise to the behavior we documented. While we cannot pinpoint the source of the variation we observed, it points to the need to explore how muscle uses a broad palette of kinematics to produce functional movement in a constantly changing environment.

## Acknowledgements

The authors gratefully acknowledge the thoughtful feedback of Joseph Powers, Abigail von Hagel and Michael Regnier on muscle biophysics; and the feedback of Valentina Staneva, Callin Switzer, Satpreet Singh and Bingni Brunton on data analysis.

## Contribution

Insert the Contribution text here.

## Data availability

Insert the Data availability text here.

## Supplementary

Insert the supplementary text text here.

