## Supplemental data for "*In vivo* x-ray diffraction and simultaneous EMG reveal the time course of myofilament lattice dilation and filament stretch"

### Supplement

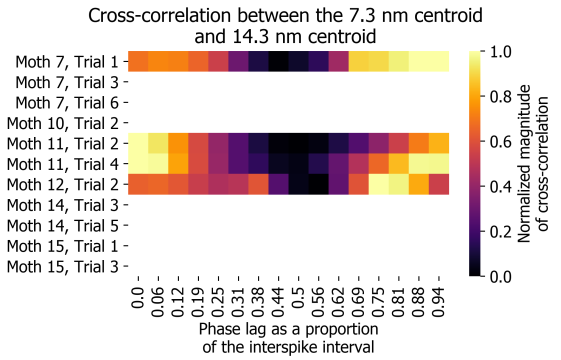


**Fig. S1. The maximum correlation between the 7.3 nm repeat and 14.3 nm repeat occurs at small but variable phase offsets across trials.** The signals are minimally correlated at a phase offset midway through the inter-spike interval (the duration of time between one muscle activation and the next). The M6 and M3 may record different structures, explaining why signals are not perfectly phase locked. The reason why this is the case follows. X-ray diffraction takes a Fourier transform of the myofilament lattice. In short exposure images the intensities recorded by the detector due to periodic structures are hard to disambiguate from noise. By averaging many images strong traces from periodic structures emerge. Frame averaging is analogous to increasing the total photon count. Similarly, reflections close to the backstop are the composite of larger numbers of photons. Signal averaging is more important when there is a low signal to noise ratio. For labile structures like cross-bridges, a large photon count is needed to create a crisp signal that rises above the background noise. In contrast the more stable thick filament backbone's periodicity is less dependent on averaging. Hence, the intensity due to the myosin crown repeat predominates close to the backstop, but as the distance from the backstop increases, the signals from labile structures will become more diffuse, while those from stable structures retain their integrity. Therefore the M6 and M3 may record different structures, explaining why signals may not be perfectly phase-locked.

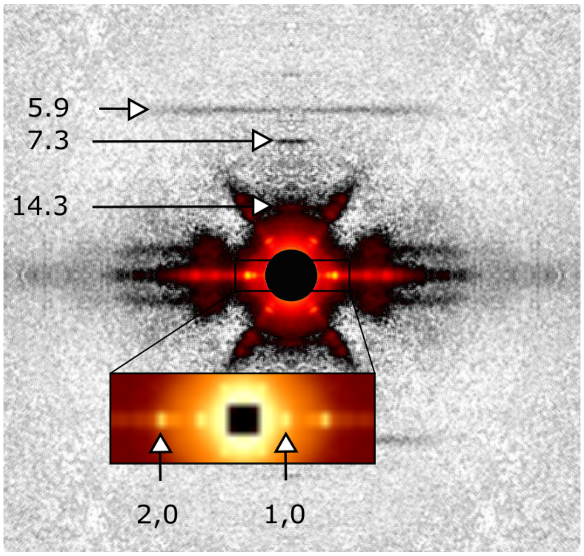


**Fig. S2. A single, x-ray diffraction image from Moth 15 trial 1.** Greater clarity could have been gained from longer exposures at the expense of reduced time resolution. Since the conversion from pixel to nanometer space by Bragg's law is non-linear, error disproportionately effects data recorded close to the backstop. An error of one pixel corresponds to 0.04 nm for the 7.3 nm spacing, while for the 5.9 actin off meridional an error of one pixel corresponds to an error of 0.03 nm. Meanwhile for the 14.3 nm reflection and the lattice spacing, which are both closer to the center of the image, an error of one pixel corresponds to 0.17 nm and 1.68 nm respectively.

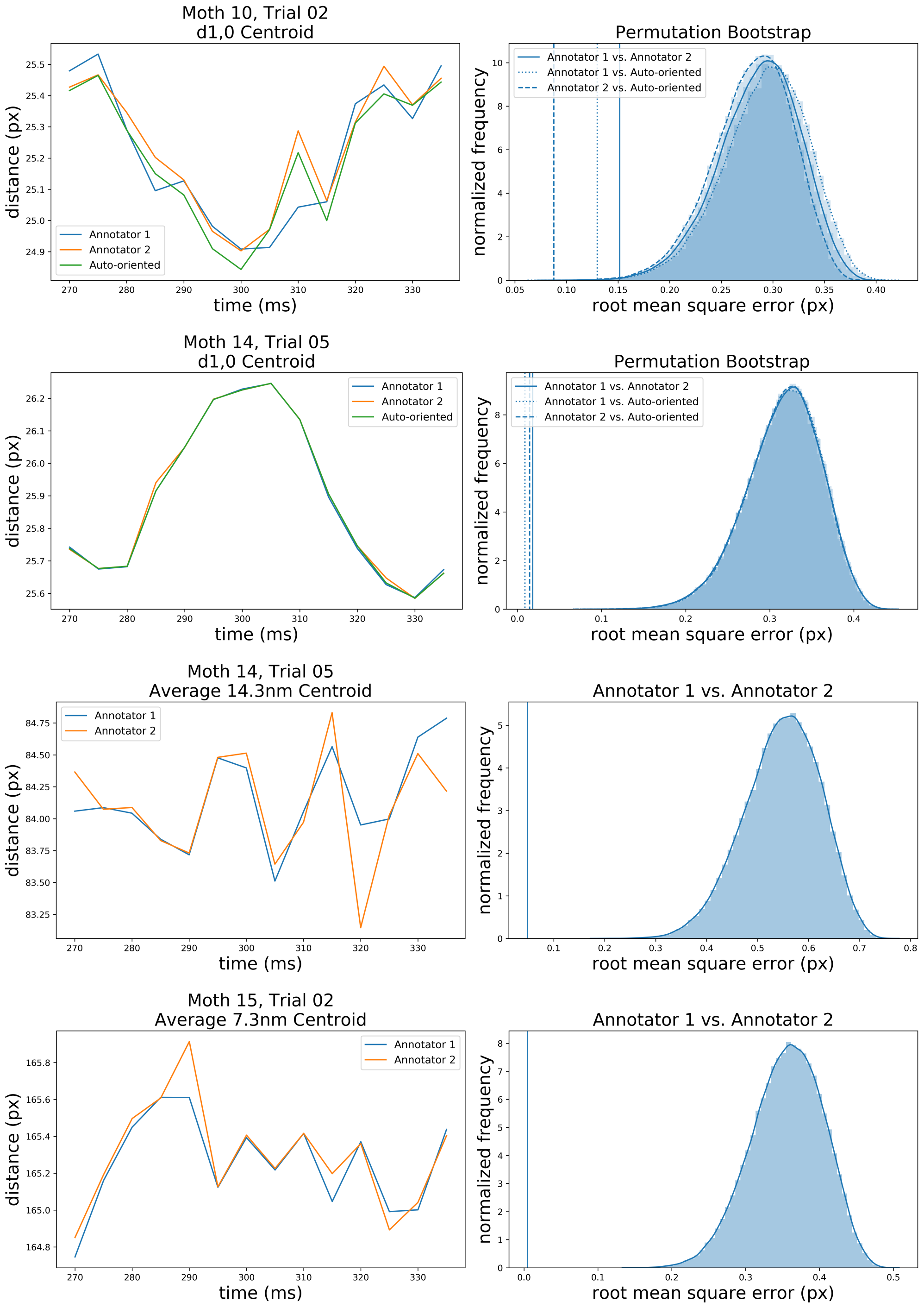


**Fig. S3. Quantification of inter-annotator error.** To verify that there was not significant inter-annotator variation we compared their pixel space annotations for a time series (a stack of images). For the d1,0 data we also compared a version of Musclex with an auto-orienting patch to Musclex. To quantify the variation between annotators shown in the time series plots on the left hand side, we computed a Root Mean Square Goodness of Fit (RMS GOF) between the two annotated image stacks. This value was compared to the distribution of RMS GOFs for 10,000 random permutations of the time series. We performed this for 4 trials with 15 time points from the d1,0 centroid, the 7.3 nm centroid and the 14.3 nm centroid. Two of the 4 cross validated trials weren't used in the final analysis; in one case because the muscle activation couldn't be determined from the EMG, and in the other case because the moth quit flying in the middle of the trial. The right hand plots show the distribution of RMS values calculated from the permutation bootstrap, with vertical lines denoting the RMS value for each annotation comparison. The annotator-comparison RMS GOF values are outside of the permutation bootstrap distribution, suggesting that the results are not explained by chance. The low annotator-comparison RMS GOF values indicate that the variation between the annotators was low. In addition to the RMS GOF, we calculated a p-value for each trial. The p-value addresses the null hypothesis that the order of the data does not affect the RMS GOF value, with a low p-value indicating that the annotations are consistent in their placement within the series. A low RMS GOF value indicates low disparity in annotator recorded values. The distributions for our data were typically unimodal. The average RMS of the d1,0 centroid was 0.07, and the average p value was 0.0008. The average RMS of the 7.3 nm was 0.70 and the average p value was 0.02. The average RMS of the 14.3 nm was 0.41, while the p value was 0.03.

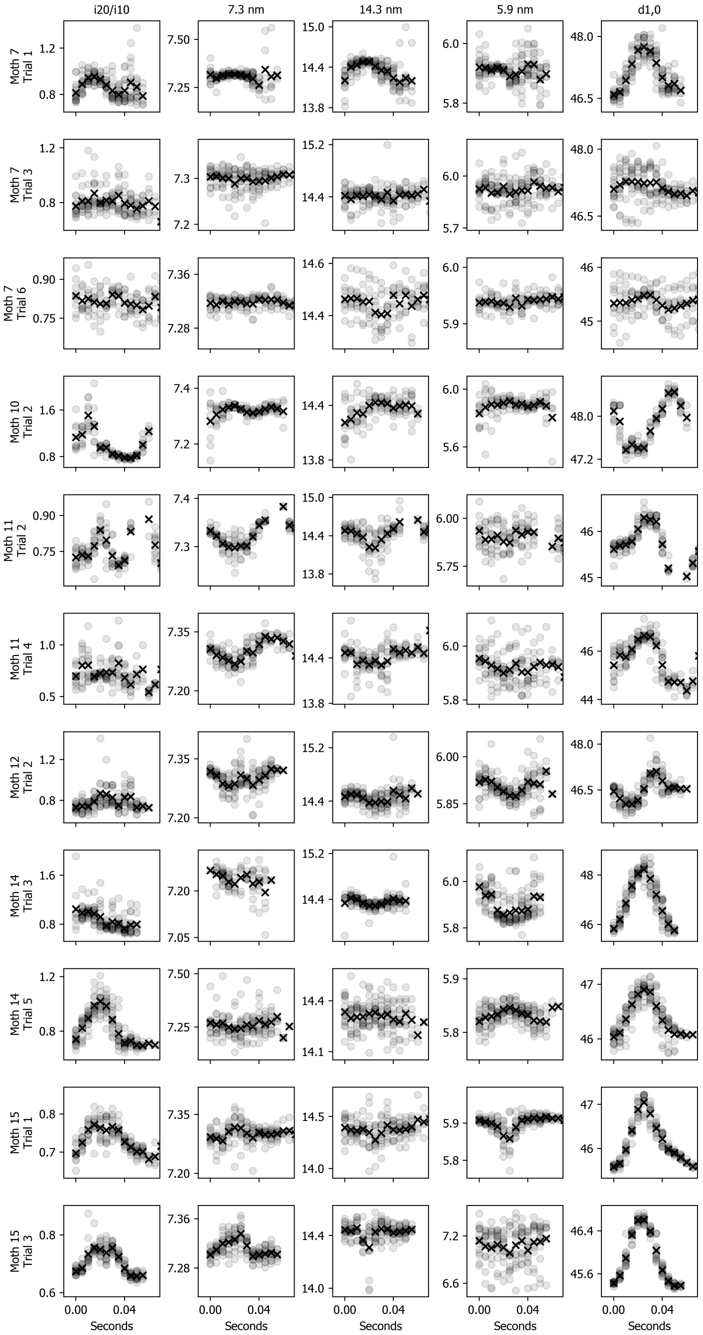


**Fig. S4. All of the data we collected is shown in this panel of STAs.**

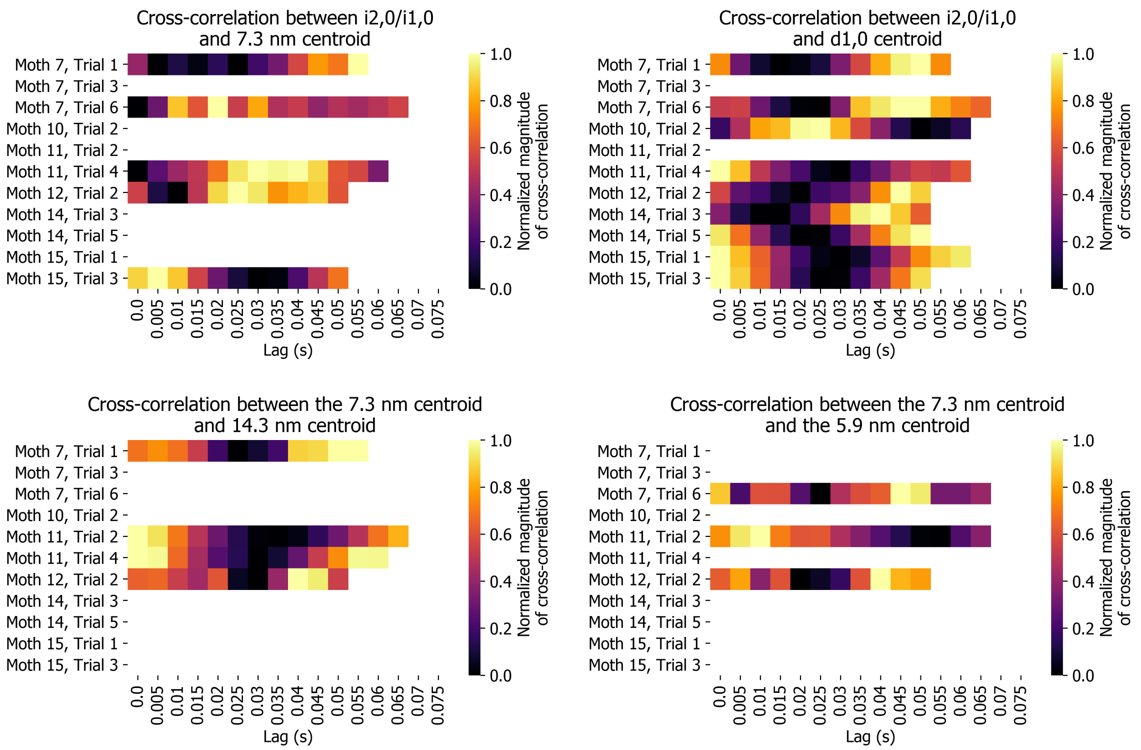


**Fig. S5. The cross-correlation between a pair of signals given an offset of the signals measured in seconds shows inconsistent phase relationships across trials**. This is in contrast to the other plots of cross correlation as a function of proportion of interspike interval. In none of these cases does a given offset in seconds yield the maximum correlation across trials.

**Table S1.** The optimal hyperparameters determined by the genetic algorithm for each trial are listed for the case where the model was trained on 75% of the data and tested on the remaining 25% of the data within a trial.

| Within trial model parameters | | | | | | | |
|---|---|---|---|---|---|---|---|
| **Trial** | **learningrate** | **nestimators** | **maxdepth** | **minchildweight** | **gamma** | **subsample** | **colsamplebytree** |
| m07_t01_15 | 0.16 | 241.0 | 7.0 | 5.1 | 0.04 | 0.77 | 0.79 |
| m07_t03_15 | 0.26 | 98.0 | 1.0 | 1.53 | 1.19 | 0.29 | 0.64 |
| m07_t06_15 | 0.05 | 135.0 | 3.0 | 0.72 | 1.22 | 0.62 | 0.34 |
| m10_t02_16 | 0.9 | 134.0 | 2.0 | 2.1 | 1.81 | 0.55 | 0.27 |
| m11_t02_16 | 0.45 | 111.0 | 9.0 | 7.38 | 2.37 | 0.59 | 0.59 |
| m11_t04_16 | 0.44 | 204.0 | 2.0 | 5.96 | 0.89 | 0.8 | 0.75 |
| m12_t02_16 | 0.43 | 98.0 | 1.0 | 7.18 | 0.79 | 0.21 | 0.42 |
| m14_t05_16 | 1.0 | 160.0 | 4.0 | 10.0 | 7.85 | 1.0 | 0.4 |
| m14_t03_16 | 0.22 | 248.0 | 8.0 | 0.16 | 0.51 | 0.62 | 0.48 |
| m15_t01_16 | 0.29 | 30.0 | 7.0 | 6.8 | 0.73 | 0.82 | 0.43 |
| m15_t03_16 | 0.34 | 48.0 | 2.0 | 6.68 | 0.01 | 0.93 | 0.23 |

**Table S2.** The optimal hyperparameters determined by the genetic algorithm for each trial are listed for the case where the model was trained on all trials except for the with-held trial upon which it was tested.

| Across trials model parameters | | | | | | | |
|---|---|---|---|---|---|---|---|
| **Trial** | **learningrate** | **nestimators** | **maxdepth** | **minchildweight** | **gamma** | **subsample** | **colsamplebytree** |
| m07_t01_15 | 0.31 | 198.0 | 5.0 | 10.0 | 7.95 | 0.85 | 0.91 |
| m07_t03_15 | 0.11 | 10.0 | 4.0 | 10.0 | 0.61 | 0.63 | 0.92 |
| m07_t06_15 | 0.05 | 26.0 | 3.0 | 6.69 | 7.42 | 0.76 | 0.13 |
| m10_t02_16 | 0.21 | 54.0 | 4.0 | 5.72 | 4.68 | 0.67 | 0.79 |
| m11_t02_16 | 0.06 | 217.0 | 1.0 | 9.41 | 7.24 | 1.0 | 0.36 |
| m11_t04_16 | 0.35 | 30.0 | 4.0 | 9.66 | 9.13 | 0.74 | 0.9 |
| m12_t02_16 | 0.07 | 19.0 | 6.0 | 6.37 | 5.45 | 0.77 | 0.4 |
| m14_t05_16 | 0.51 | 319.0 | 7.0 | 10.0 | 7.39 | 0.62 | 0.49 |
| m14_t03_16 | 0.59 | 55.0 | 7.0 | 5.18 | 2.58 | 0.99 | 0.88 |
| m15_t01_16 | 0.09 | 136.0 | 5.0 | 10.0 | 8.6 | 0.21 | 0.52 |
| m15_t03_16 | 0.25 | 179.0 | 10.0 | 10.0 | 6.04 | 0.12 | 0.12 |
